# ribosomal DNA instability as a potential cause of karyotype evolution

**DOI:** 10.1101/2022.07.10.499502

**Authors:** Duojia Li, Dhyey Gandhi, Tomohiro Kumon, Yukiko M. Yamashita

## Abstract

Karyotype refers to the configuration of the genome into a set of chromosomes. The karyotype difference between species is expected to impede various biological processes, such as chromosome segregation or meiotic chromosome pairing, potentially contributing to incompatibility. Karyotypes can rapidly change between closely related species and even among populations of the same species. However, the forces driving karyotype evolution are poorly understood. Here we describe a unique karyotype of a *D. melanogaster* strain isolated from the Seychelles archipelago. This strain has lost the ribosomal DNA (rDNA) locus on the X chromosome. Because the Y chromosome is the only other rDNA-bearing chromosome, all females carry at least one Y chromosome as the source of rDNA. Interestingly, we found that the strain also carries a truncated Y chromosome (Y^S^) that is stably maintained in the population despite its inability to support male fertility. Our modeling and cytological analysis suggest that the Y chromosome has a larger negative impact on female fitness than the Y^S^ chromosome. Moreover, we generated an independent strain that lacks X rDNA and has a karyotype of XXY females and XY males. This strain quickly evolved multiple karyotypes: two new truncated Y chromosomes (similar to Y^S^), as well as two independent X chromosome fusions that contain the Y-derived rDNA fragment, eliminating females’ dependence on the Y chromosome. Considering that Robertsonian fusions frequently occur at rDNA loci in humans, we propose that rDNA loci instability is a driving force of karyotype evolution.

## Introduction

Karyotype refers to the configuration of the genome into a set of chromosomes. Chromosomes are the physical units that carry genomic information, and are subjected to cell biological constraints such as mitotic chromosome segregation and meiotic chromosome pairing. Accordingly, different karyotypes can be incompatible with each other even if they carry the same genetic information, leading to speciation (Ayala and Coluzzi, 2005; King, 1995). For example, Indian muntjac deer and Chinese muntjac deer are close enough to produce viable hybrids, however, these hybrids are sterile (Liming and Pathak, 1981). The Indian muntjac contains only six (female) or seven (male) chromosomes whereas the Chinese muntjac contains 46 chromosomes. This drastic karyotype difference may cause incompatibility during meiotic chromosome pairing, potentially explaining hybrid sterility, although the detailed underlying cell biological causes of their sterility are not well understood. Karyotype differences can have a profound negative impact on fitness even between individuals within the same species. Two telocentric chromosomes fuse to form a Robertsonian translocation in approximately 1 in every 1000 newborn humans (Hamerton et al., 1975), and a meiotic trivalent of these three chromosomes (i.e. the Robertsonian fusion chromosome and the two telocentric chromosomes) is known to cause meiotic segregation errors, leading to subfertility (de Villena and Sapienza, 2001). These observations indicate that karyotype difference, despite the compatibility of the information encoded in the genome, is sufficient to cause a fitness cost. However, the processes and driving forces of karyotype evolution leading to speciation are poorly understood.

Ribosomal DNA (rDNA), a genomic element required for ribosomal biogenesis, exists as tandemly repeated units of rRNA genes in eukaryotic genomes, often divided into multiple rDNA loci, each containing hundreds of rDNA copies (McStay, 2016). For example, the *D. melanogaster* genome has two rDNA loci, one on the X chromosome and one on the Y chromosome (Figure 1A). The human genome has five rDNA loci per haploid genome, on the autosomes, whereas the mouse genome has six rDNA loci per haploid genome, also on the autosomes (McStay, 2016).

**Figure 1.**
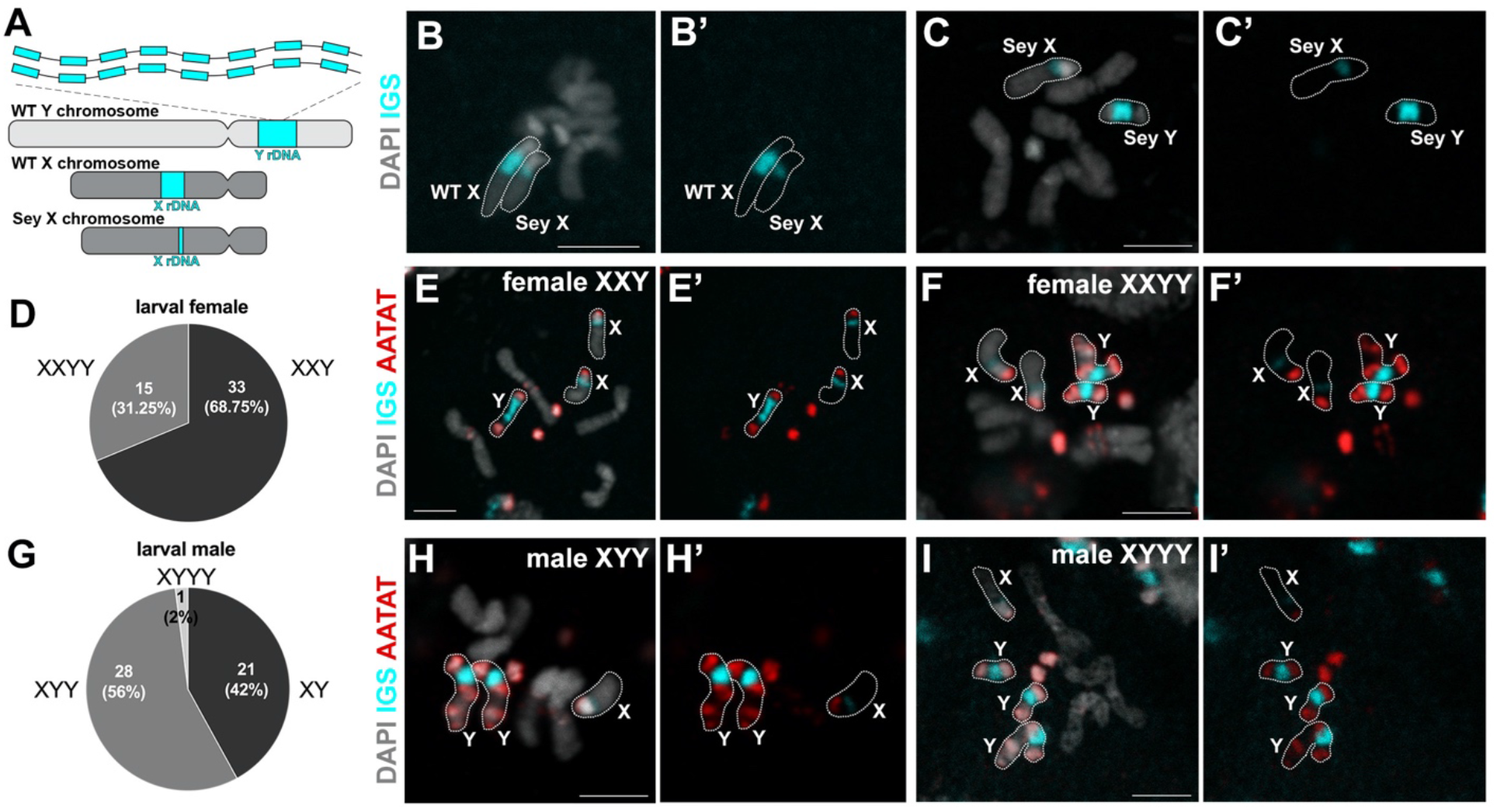
*D. mel*^*Sey*^ strain has minimal rDNA on X, necessitating females to carry Y chromosomes. A) Schematics of wild type (WT) X, Y chromosomes and *D. mel*^*Sey*^ X chromosome (note that *D. mel*^*Sey*^ Y was indistinguishable from WT Y with respect to rDNA by FISH). B) DNA FISH on the mitotic chromosome spread of larval neuroblasts carrying wild type (Oregon-R) X and *D. mel*^*Sey*^ X. Intergenic spacer (IGS) of rDNA was used to probe for rDNA loci, AATAT satellite DNA was used to aid the identification of chromosomes. Bar: 3μm in all panels. C) DNA FISH on the mitotic chromosome spread of larval neuroblasts of *D. mel*^*Sey*^ XY male. D) Distribution of *D. mel*^*Sey*^ female karyotype assessed by DNA in situ hybridization on the mitotic chromosome spread of larval neuroblasts. E) DNA FISH on the mitotic chromosome spread of larval neuroblasts from a *D. mel*^*Sey*^ XXY female. F) DNA FISH on the mitotic chromosome spread of larval neuroblasts from a *D. mel*^*Sey*^ XXYY female. G) Distribution of *D. mel*^*Sey*^ male karyotype assessed by DNA in situ hybridization on the mitotic chromosome spread of larval neuroblasts. H) DNA FISH on the mitotic chromosome spread of larval neuroblasts from *D. mel*^*Sey*^ XYY male. I) DNA FISH on the mitotic chromosome spread of larval neuroblasts from *D. mel*^*Sey*^ XYYY male.

Interestingly, the positions of rDNA loci are not well conserved, even between species with nearly syntenic genome arrangement (Cazaux et al., 2011; Mazzoleni et al., 2018; Roy et al., 2005; Sochorová et al., 2017). Moreover, within-species variations in the location of rDNA loci have been reported in multiple species of animals and plants (Baicharoen et al., 2016; Hwang et al., 2011). These observations suggest that rDNA loci are a relatively unstable region of the genome. This is not too surprising, given that repetitive loci are prone to intrachromatid recombination, which has been shown to reduce rDNA copy number (Nelson et al., 2019). Although different mechanisms can help recover rDNA copy number (Hawley and Tartof, 1985), a locus may be lost at a low but significant frequency in an evolutionary time scale. For example, whereas *D. melanogaster* has rDNA loci on X and Y chromosomes, its closely related species, *D. simulans* and *D. sechellia* (Roy et al., 2005), have lost a functional rDNA locus from the Y chromosome, suggesting the instability of rDNA locus. Moreover, homologous recombination between rDNA loci on non-homologous chromosomes would cause chromosome rearrangements, leading to changes in karyotype. Indeed, the flanking regions of multiple rDNA loci have conserved DNA sequences, indicating a history of DNA recombination in the creation of the rDNA loci (Floutsakou et al., 2013). Despite these observations, it remains unknown whether and how the instability of rDNA may contribute to karyotype evolution.

Here, we report a *D. melanogaster* strain isolated from the Seychelles archipelago (referred to as *D. mel*^Sey^ hereafter) that has lost rDNA from the X chromosome, necessitating females to carry a Y chromosome as the sole source of rDNA. Interestingly, we found that *D. mel*^Sey^ carries a truncated Y chromosome (Y^S^, carrying rDNA but losing a large part of the long arm of the Y chromosome) in addition to the normal, full length Y chromosome. Combining cytological analysis and mathematical modeling, our study suggests that Y^S^ is well-maintained in the population, despite its inability to support male fertility, because Y^S^ imposes a less negative impact on female fitness compared to the full-length Y chromosome, which is known to reduce female lifespan due to an excess of heterochromatin (Branco et al., 2017; Brown et al., 2020a; 2020b). Moreover, we constructed an independent laboratory stock of *D. melanogaster* that lacks X rDNA, forcing all individuals to carry Y chromosome as the source of rDNA. Intriguingly, multiple new chromosomes rapidly evolved in this stock: two new truncated Y chromosomes similar to Y^S^, two new X chromosomes (X* and X**). The X* and X** chromosomes are fusions between the original X chromosome lacking rDNA and an rDNA-containing piece of the Y chromosome. Thus, females with X* or X** no longer carry the full-length Y chromosome. These results suggest that the necessity and instability of rDNA, combined with the well-established burden of the Y chromosome in females (Branco et al., 2017; Brown et al., 2020a; 2020b), is a potent driver of karyotype changes. Considering that Robertsonian fusions frequently occur at rDNA loci in humans, we propose that rDNA loci instability is a universal and major driving force of karyotype evolution.

## Results

### *D. melanogaster* Seychelles strain has a minimal amount of rDNA on X, leading to an unusual karyotype

The Seychelles strain of *D. melanogaster* (*D. mel*^Sey^) was isolated from the Seychelles archipelago in 1987. In the course of investigating sequence variations of rDNA in *D. melanogaster* strains, we unexpectedly found that *D. mel*^Sey^ contains an insufficient amount of rDNA on their X chromosome. Specifically, genetic crosses between *D. mel*^Sey^ and a *D. melanogaster* strain carrying a complete rDNA deletion on the X chromosome (*bb*^*158*^) revealed that female progenies (X_Seychelles_/*bb*^*158*^) are inviable. To investigate rDNA copy number, we performed DNA fluorescence in situ hybridization (FISH) on mitotic chromosomes of larval neuroblasts using the intergenic spacer of rDNA (IGS) probe. We observed a greatly reduced rDNA locus on the *D. mel*^Sey^ X chromosome compared to the wild type (Oregon-R) X chromosome (Figure 1B), or to the Y chromosome of *D. mel*^Sey^ (Figure 1C). We infer that rDNA on the *D. mel*^Sey^ X chromosome is insufficient to support female viability even when present in two copies (X_Seychelles_/X_Seychelles_). Accordingly, we found that all females of the *D. mel*^Sey^ strain contained at least one Y chromosome (100%, n=48 females examined): 68.75% of females were XXY (Figure 1D, E), whereas the remaining females (31.25%) were XXYY (Figure 1D, F), with the Y chromosome serving as the source of rDNA. It should be noted that *Drosophila* sex determination is based on the X chromosome: autosome ratio, not the presence of the Y chromosome. The Y chromosome contains an rDNA locus, as well as genes required for male fertility. Although rDNA on a single Y chromosome was sufficient for *D. mel*^Sey^ viability, many individuals (31.25% of females and 58% of males) carried more than one Y chromosome (Figure D, F-I), which is likely caused by transmission of Y chromosomes from both parents.

These results revealed that *D. mel*^Sey^ has a unique karyotype, with both males and females carrying Y chromosomes, likely due to the reduction of rDNA on the X chromosomes.

### *D. mel*^Sey^ carries a new, truncated Y^S^ chromosome that cannot support male fertility

We noticed that some of the *D. mel*^Sey^ Y chromosomes appeared to be shorter than others (Figure 1F, I). We performed DNA FISH with various Y chromosome satellite DNA probes, and confirmed the presence of a truncated Y chromosome (Y^S^) lacking the distal half of the long arm of the Y chromosome (Figure 2A-D, Supplementary Figure 1).

**Figure 2.**
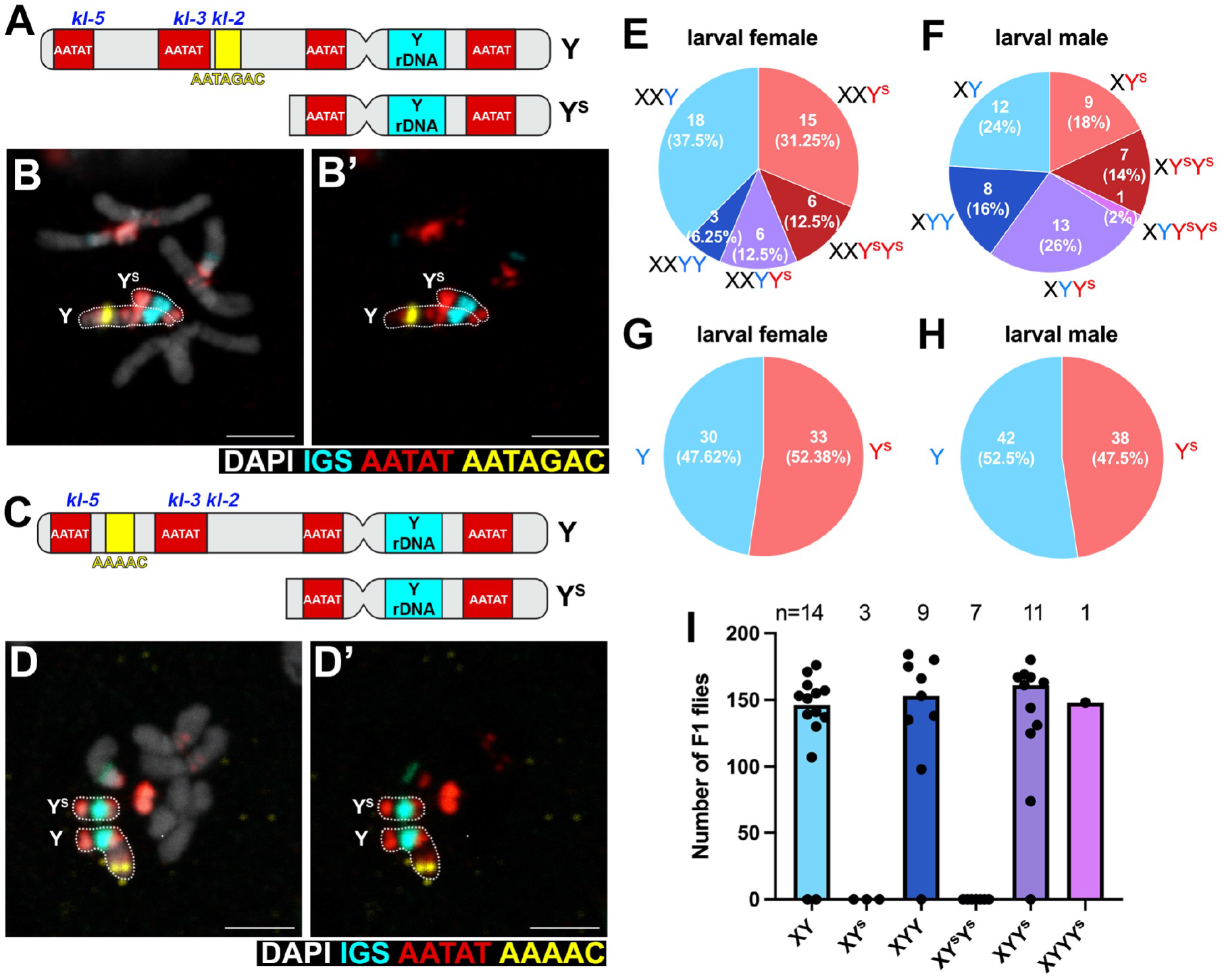
*D. mel*^*Sey*^ strain carries full-length Y and truncated Y (Y^S^) chromosomes. A) Location of satellite DNA (AATAT, AATAGAC) used in DNA FISH in panel B to distinguish Y and Y^S^. The locations of male fertility factors (*kl-2. -3, -5*) deleted in the Y^S^ chromosome are indicated. B) DNA FISH on larval neuroblast mitotic chromosome spread showing AATAGAC satellite is present on Y but not Y^S^. Bar: 3 μm in all panels. C) Location of satellite DNA (AATAT, AAAAC) used in DNA FISH in panel D to distinguish Y and Y^S^. D) DNA FISH on larval neuroblast mitotic chromosome spread showing AAAAC satellite is present on Y but not Y^S^. E) Frequencies of female karyotypes. n=48 larval females examined by DNA FISH. F) Frequencies of male karyotypes. n=50 larval males examined by DNA FISH. G) Frequencies of Y vs. Y^S^ chromosomes in 48 larval females. H) Frequencies of Y vs. Y^S^ chromosomes in 50 larval males. I) Fertility of *D. mel*^*Sey*^ males of the indicated karyotypes.

We examined the frequencies of Y and Y^S^ chromosomes in the population by karyotyping individual flies using DNA FISH on mitotic chromosome spreads of larval neuroblasts (Figure 2E, F). These data revealed that both males and females carry Y and Y^S^ chromosomes at similar frequencies: 37.5% of *D. mel*^Sey^ females were XXY and 31.25% were XXY^S^, with XXYY, XXY^S^Y^S^, XXYY^S^ representing smaller populations (6.25%, 12.5%, 12.5%, respectively, n=48 females examined, Figure 2E). In males, 24% were XY, 18% were XY^S^, 16% were XYY, 14% were XY^S^Y^S^, and 26% were XYY^S^ (n=50 males examined, Figure 2F). Overall, Y and Y^S^ chromosomes were present at similar frequencies in both males and females: among total Y chromosomes, Y: Y^S^ was 47.62%: 52.38% in females (n=63 Y or Y^S^ chromosomes in 48 females), and 52.5%: 47.5% in males (n=80 Y or Y^S^ chromosomes in 50 males) (Figure 2G-H).

The chromosomal region that was deleted in the Y^S^ chromosome contains some fertility factors, including *kl-5, kl-3* and *kl-2*, which encode axonemal dynein subunits that are essential for sperm development (Carvalho et al. 2000) (Figure 2A, C, Supplementary Figure 1). Therefore, we hypothesized that males that carry only the Y^S^ chromosome are sterile. To test this possibility, we mated individual males with 3 wild type (*yw*) females, and after a week, individual males were subjected to karyotyping using ddPCR, which can precisely measure the copy number of the genes of interest (Method). We indeed found that males that contain only Y^S^ chromosomes (XY^S^ and XY^S^Y^S^) were sterile, whereas males with at least one full length Y chromosome were fertile (Figure 2I).

These results reveal a complex karyotype in *D. mel*^Sey^, where two kinds of Y chromosomes are present. Male fertility requires a full-length Y, whereas male and female viability require a full-length Y or truncated Y^S^.

### Karyotype stability modeling predicts that Y has a higher fitness cost than Y^s^ in females

Considering that the Y^S^ chromosome cannot support male fertility, it is striking that the Y^S^ chromosome is so common within the population: nearly half of the total Y chromosomes were the Y^S^ in both males and females. We simulated the dynamics of Y and Y^S^ chromosome frequencies in the population over 200 generations, and our modeling suggested that the Y^S^ chromosome would quickly disappear from the population if it does not provide any benefit, given that it is detrimental to male fertility (Figure 3A).

**Figure 3.**
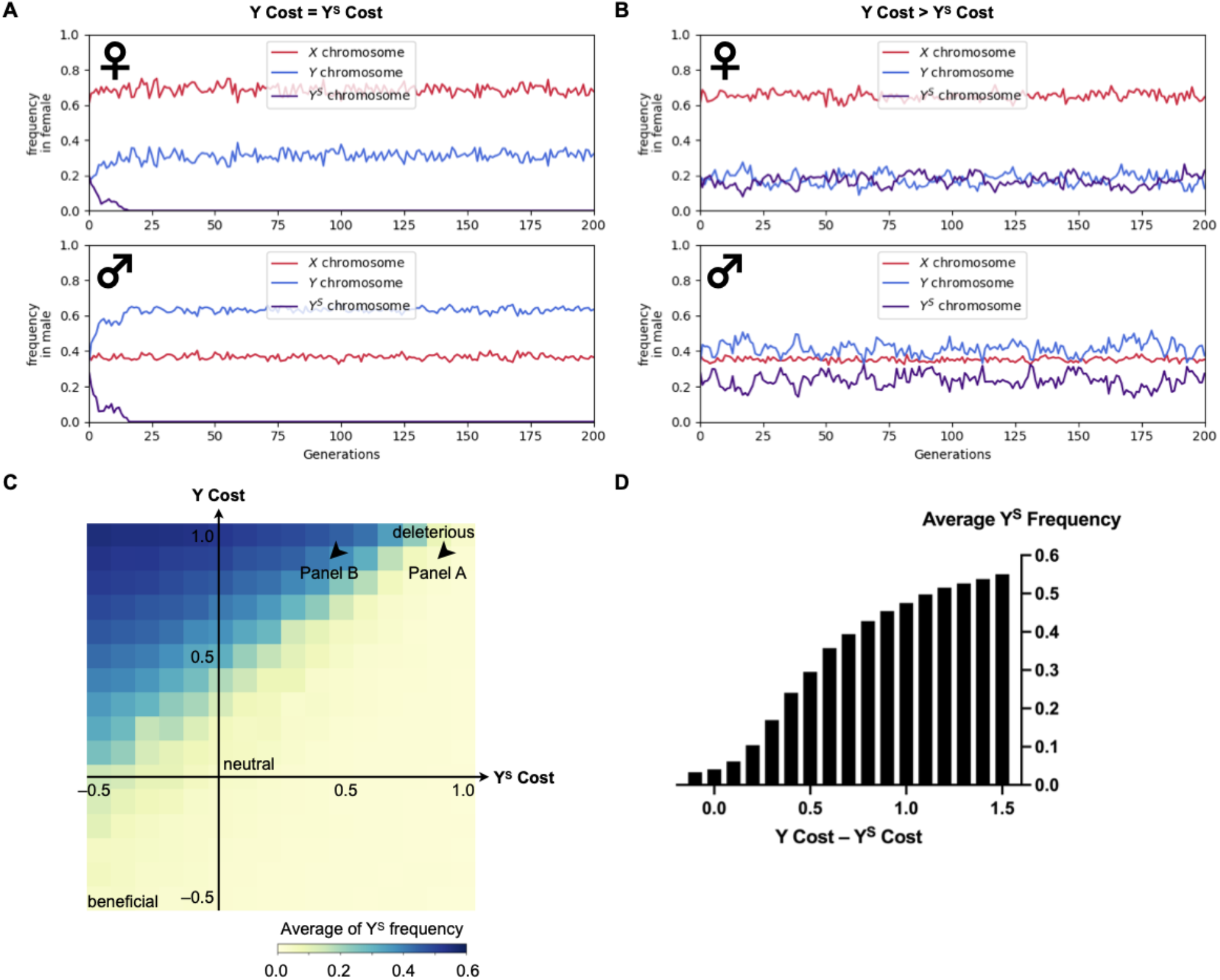
The fitness difference of Y and Y^S^ chromosomes in females affects the Y^S^ frequency in the population. A) Sex chromosome frequencies over two hundred generations when the female cost of the Y chromosome is equal to that of the Y^S^ chromosome. The Y^s^ chromosome rapidly disappears from the population. B) Sex chromosome frequencies over two hundred generations when the female cost of the Y chromosome is twice as much as that of the Y^S^ chromosome (the exact combination of Y/Ys costs is plotted in panel C). Top panel shows sex chromosome frequencies in females and bottom panel shows those in males. C) The average Y^S^ frequency over a hundred generations with varying female costs of Y and Y^S^ chromosomes. In panels B and C, the average Y^S^ frequency is the percentage of Y^S^ chromosomes among all the Y chromosomes. Briefly, the chromosome is regarded as deleterious (making females infertile) when the female cost is 1, neutral when the cost is 0, and beneficial when the cost has negative values (Method). Arrowheads show the parameters used to simulate Panels A and B. The values of “Y Cost” and “Y^S^ Cost” are equal to the values of *y* and *s* in Method, respectively. D) The fitness differences of Y and Y^S^ chromosomes and corresponding average Y^S^ frequencies. When the fitness differences become small, the average Y^S^ frequency decreases and often goes extinct from the population.

It has been reported that, in females with a Y chromosome, the extra heterochromatin dilutes heterochromatin marks throughout the entire genome, leading to changes in gene expression patterns, especially in the ovaries (Branco et al., 2017). Moreover, lifespan is inversely correlated with the number of Y chromosomes (Brown et al., 2020a; 2020b). Therefore, we hypothesized that a higher fitness cost of the full-length Y compared to the Y^S^ chromosome in females can potentially explain the persistence of the Y^S^ chromosome in the *D. mel*^Sey^ strain. We modeled this possibility by simulating the dynamics of Y and Y^S^ chromosome frequencies in the population over 200 generations under different female costs of the Y vs. Y^S^ chromosomes (Method and Supplementary Figure 2). Our modeling predicts that, for example, if the female cost of the Y chromosome is two times higher than the Y^S^ chromosome, then both Y and Y^S^ chromosomes are stably maintained in the population for hundreds of generations (Figure 3B). The heatmap of Figure 3C shows the average frequency of the Y^S^ chromosome over a hundred generations under different female costs of the Y vs. Y^S^ chromosomes. For example, when the cost of the Y chromosome minus the cost of the Y^S^ chromosome is larger than 0.5, the probability of the Y^S^ persistence becomes more than 90%. On the contrary, our simulation suggests that the Y^S^ chromosome reduces its frequency and often becomes extinct as the fitness difference of Y and Y^S^ chromosomes becomes small (Figure 3D). Taken together, these modeling results suggest that the Y^S^ chromosome is maintained because it is less costly than the Y chromosome in females, under the unique circumstance that all females must carry Y chromosome as a source of rDNA.

### *D. mel*^Sey^ XXY^S^ females have less severe ovariole degeneration than *D. mel*^Sey^ XXY females

To characterize a potential higher fitness cost of the Y compared to Y^S^ chromosome in females, we examined females of different karyotypes. The frequencies of *D. mel*^Sey^ XXY vs. XXY^S^ females were similar in 3rd instar larvae (Figure 2E) and adult (not shown), suggesting that Y^S^ does not improve female viability during development compared to Y. To examine female fertility, we crossed individual females of the *D. mel*^Sey^ strain to three wild type (*yw*) XY males and after a week, parents were removed, and the female was subjected to karyotyping using ddPCR, as described above. The number of progeny from the week of mating was determined when they eclose as adult flies. We did not observe significant differences in the fertility of XXY vs. XXY^S^ females during the first week (Figure 4A). However, we noted that females with multiple Y or Y^S^ chromosomes were significantly less fertile than females with only one Y or Y^S^ chromosome (Figure 4A), suggesting that the Y chromosome poses dose-sensitive negative impacts on female fertility.

**Figure 4.**
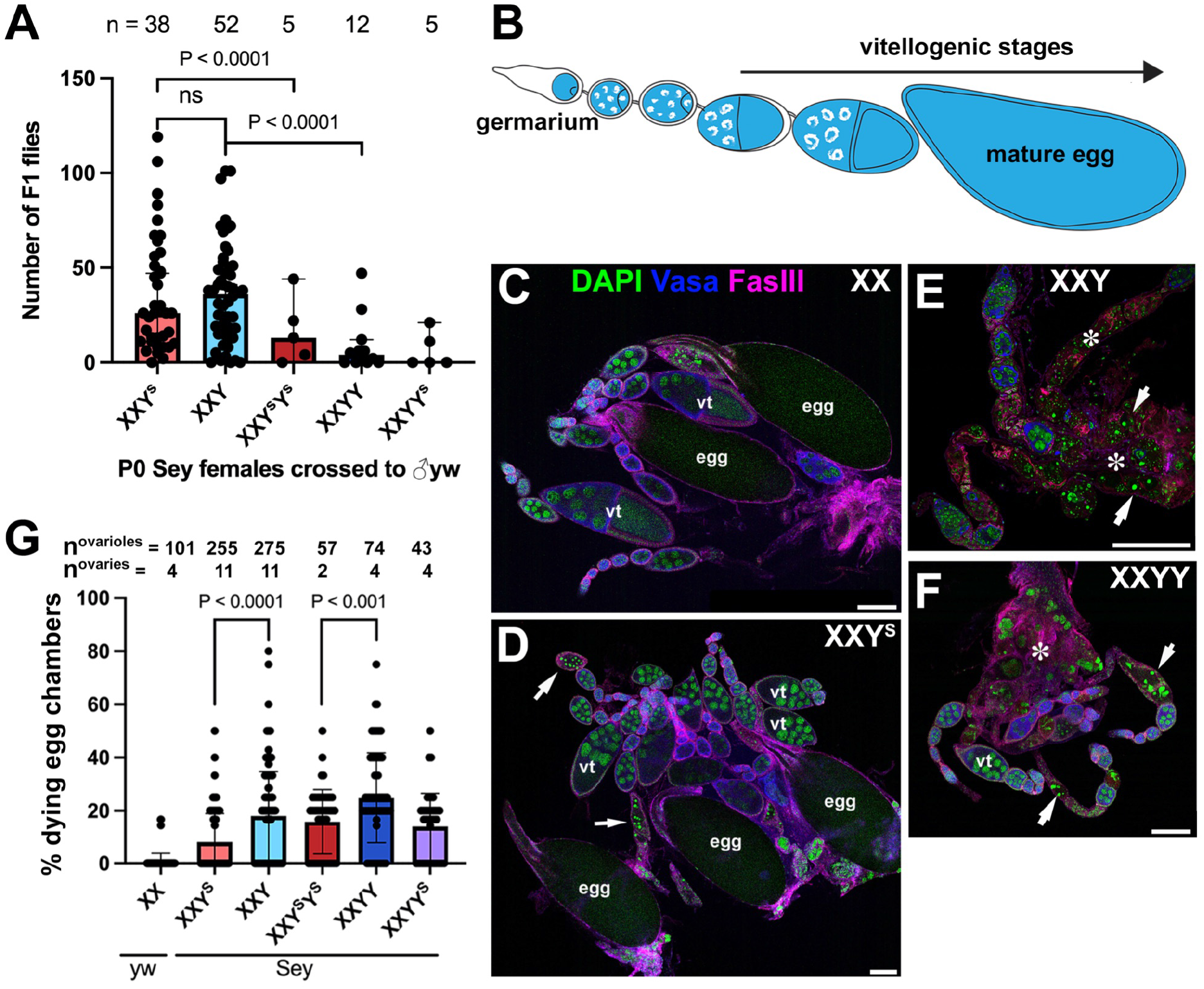
Y^S^ is less detrimental to oogenesis than the full-length Y chromosome. A) Fertility of females of the indicated karyotypes, showing that having 2 Y or Y^S^ chromosomes reduces female fertility. p-values were calculated using an unpaired Student’s t-test with Welch’s correction (assuming unequal variances) with n representing the number of P0 individual males. B) Diagram of the Drosophila ovariole. C-F) Representative immunofluorescent images of ovarioles in (C) wild type (*yw*) XX, (D) *D. mel*^*Sey*^ XXY^S^, (E) *D. mel*^*Sey*^ XXY, and (F) *D. mel*^*Sey*^ XXY females stained for Vasa (germ cells, blue) and Fas III (cell membranes, magenta), counterstained with DAPI (green). Arrows indicate dying egg chambers. Asterisks (*) indicate degenerated egg chambers. Representative egg chambers are marked with ‘egg’ (mature eggs) and ‘vt’ (vitellogenic stages). Bar: 100 μm. G) Frequency of ovarioles with dying egg chambers in the indicated karyotypes. p-values were calculated using an unpaired Student’s t-test with Welch’s correction (assuming unequal variances) with n representing the number of ovarioles.

To investigate the impact of Y and Y^S^ chromosomes on oogenesis, we performed a cytological examination of ovaries from various karyotypes of *D. mel*^Sey^ females. The ovariole is a unit of egg production, where developing egg chambers of increasingly advanced stages are connected in a string: oogenesis starts at the germarium, leading to the production of mature eggs at the other end (Figure 4B). A checkpoint mechanism culls defective egg chambers at the beginning of the vitellogenic stage, when the transfer of cytoplasmic contents from nurse cells to the oocytes initiates (Peterson et al. 2015).

Ovarioles in wild type (*yw*) XX females rarely contained dying egg chambers, which were detected by their highly condensed DNA (Figure 4C, G). Ovarioles from *D. mel*^Sey^ XXY^S^ females were also relatively healthy, with many egg chambers containing mature eggs, although there was a slight increase in the frequency of dying egg chambers compared to wild type (Figure 4D, G). In contrast, *D. mel*^Sey^ XXY and XXYY females had considerably more dying egg chambers than XXY^S^ (Figure 4E-G, arrows), and the ovarioles were frequently degenerated after the vitellogenic stage (Figure 4E, F, asterisks). Taken together, these results suggest that the Y^S^ chromosome is less detrimental than the Y chromosome for oogenesis. However, the difference between oogenesis in females with the Y vs. Y^S^ chromosomes was rather mild (Figure 4G). Therefore, we speculate that there are additional aspects of female fitness that make Y^S^ favored to allow persistence of this ‘male sterile’ chromosome. Given the reported reduction in lifespan imposed by derepression of Y-derived heterochromatin (Branco et al., 2017; Brown et al., 2020a; 2020b), it is possible that *D. mel*^Sey^ XXY^S^ females have a better lifetime fertility compared to *D. mel*^Sey^ XXY females: however, the need for individual karyotyping by ddPCR in our fertility assay precluded us from examining female fertility beyond the first week, and thus we could not assess lifetime fertility.

### Adaptive karyotype changes in females carrying the full-length Y chromosome

Although we speculate that the Y^S^ chromosome of the *D. mel*^Sey^ strain may be a result of adaptation to reduce the burden of Y chromosome on female reproduction, it is difficult to draw a conclusion based on one unique incidence discovered in this strain. To further explore how females respond to the potential burden of the Y chromosome, we independently constructed a strain in which females must carry the full-length Y chromosome. Briefly, the *bb*^*158*^ X chromosome entirely lacks rDNA (Figure 5A) and can be maintained with a balancer X chromosome (e.g., FM6) that carries rDNA. In this strain, females are typically *bb*^*158*^/FM6 or FM6/FM6, and males are *bb*^*158*^/Y or FM6/Y. However, similar to *D. mel*^Sey^, a small fraction of females is *bb*^*158*^/*bb*^*158*^/Y and rely on the rDNA on the Y chromosome for survival. A strain without X rDNA was generated by selecting virgin *bb*^*158*^/*bb*^*158*^/Y females and *bb*^*158*^*/Y* males. We passaged this stock (‘*bb*^*158*^ stock’, hereafter) in separate lineages without any selection for multiple generations (Figure 5B). Note that some individuals carried multiple Y chromosomes, similar to the *D. mel*^Sey^ strain, likely due to biparental transmission of the Y chromosomes.

**Figure 5.**
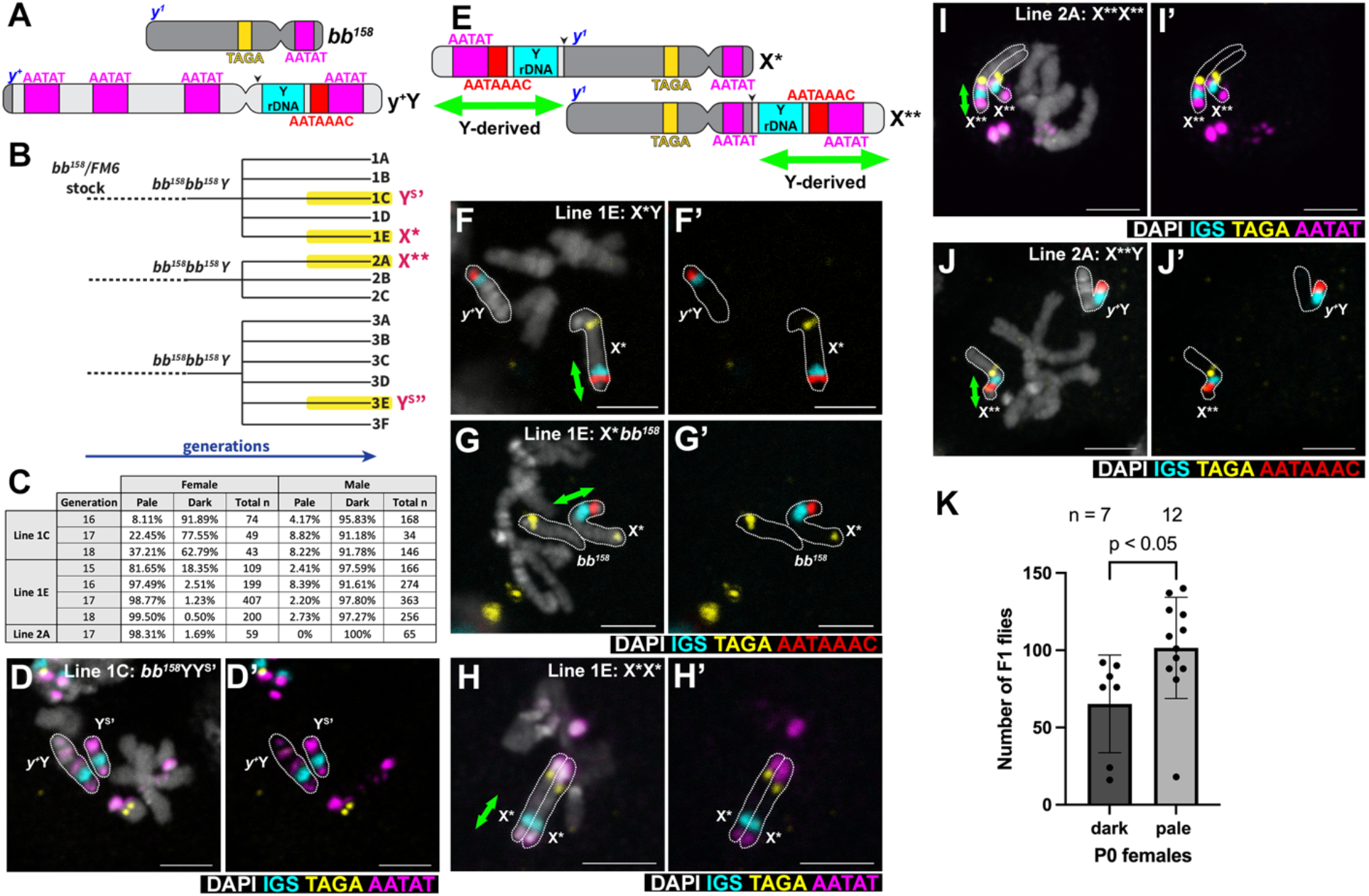
An X chromosome fusion with Y rDNA emerges to replace the full length Y chromosome in the *bb*^*158*^/*bb*^*158*^/*Y* stock. A) Schematics of *bb*^*158*^ (lacking rDNA) X chromosome with *yellow*^*1*^ mutation (*y*^*1*^) and *y*^*+*^Y chromosome marked by *yellow+* (*y*^*+*^). B) Lineage tree of derivative lines from *bb*^*158*^/*bb*^*158*^/Y stock. Three independent *bb*^*158*^/*bb*^*158*^/Y lineages were derived from the *bb*^*158*^/FM6 stock, each of which was split into several sub lineages (e.g. 1A through 1E) before the emergence of pale body-colored animals. C) Frequency of pale vs. dark body-colored females and males in the indicated lines. D) DNA FISH on a larval neuroblast mitotic chromosome spread from a *bb*^*158*^/*y*^*+*^Y/Y^S^*’* male. Bar: 3 μm in all panels. E) Schematic of X* and X** chromosomes that carry a part of the Y chromosome short arm containing rDNA. Black arrow points to the inferred breakpoints. Green double arrows mark the part of X* or X** originated from Y. F-J) DNA FISH on a larval neuroblast mitotic chromosome spread from an (F) X*/*y*^*+*^Y male; (G) *bb*^*158*^/X* female; (H) X*/X* female; (I) X**/X** female; and (J) X**/ *y*^*+*^Y male. K) Fertility assay of dark vs. pale body-colored female from the lineage 1E. p-values were calculated using an unpaired Student’s t-test with Welch’s correction (assuming unequal variances) with n representing the number of P0 females.

In this stock, the Y chromosome carries a marker, *yellow*^*+*^ (*y*^*+*^), that confers the dark body color of the wild type, while the *bb*^*158*^ chromosome has the recessive *yellow*^*1*^ (*y*^*1*^) mutation that confers the pale body color when homozygous (Figure 5A). Therefore, *bb*^*158*^/*bb*^*158*^/*y*^*+*^Y females and *bb*^*158*^/*y*^*+*^Y males have a dark body color. Intriguingly, we noticed that a small number of females had a pale body color after multiple generations in four of the lineages (Figure 5B), suggesting that they have likely lost *y*^*+*^ and no longer carry the full-length Y chromosome.

In two separate lineages (denoted lineage 1C and lineage 3E), pale females persisted in the population but never become dominant in either sex (Figure 5B, C), Using DNA FISH with various satellite DNA probes, we found that lineage 1C contains a truncated Y chromosome (Y^S^’) that carries the rDNA, resembling but somewhat shorter than Y^S^ of *D. mel*^Sey^ (Figure 5D). Lineage 3E (Y^S^’’) also contained a similar Y chromosome truncation (Supplementary Figure 3). These truncated Y chromosomes lost the *y*^*+*^ gene that is located near the telomeric end of the long arm (Figure 5A, *y*^*+*^Y chromosome), therefore females carrying only this truncated Y chromosome would be pale colored. However, similar to *D. mel*^Sey^ strain, this chromosome cannot support male fertility, likely limiting the propagation of pale colored animals in the population.

In another lineage (denoted lineage 1E) most females became pale body colored (81.65%, n=109 females scored), whereas the males mostly had dark body color (97.59%, n=166 males scored) by generation 15 (Figure 5B, C). From generations 16 to 18, the frequency of pale females increased further, reaching at 99.5%, whereas males still mostly had dark body color (Figure 5C). DNA FISH analysis of this lineage revealed that females with the pale body color no longer carry the full-length Y chromosome but instead carry an X chromosome fusion (denoted X*), in which a part of the Y chromosome containing the rDNA was fused to the long arm of the X chromosome (Figure 5E-H). DNA FISH with the Y short arm-specific satellite DNA (AATAAAC) further confirmed that the rDNA fragment attached to the X* chromosome indeed originated from the Y chromosome (Figure 5F, G). The AATAT locus next to the Y centromere on the Y long arm is not included in X* (Figure 5H), suggesting that the breakpoint on the Y chromosome was likely between the Y rDNA and the Y centromere. In addition, the rDNA (IGS) signal intensity is comparable between Y and X*, suggesting that X* contains most of Y rDNA (Figure 5F). X*/X* females are viable, and the X* chromosome morphology resembles the *bb*^*158*^ chromosome (Figure 5G). We infer that X* does not lack any essential genes on the long arm of the X chromosome and likely contains a (near) full-length *bb*^*158*^ chromosome.

In another lineage (denoted 2A), pale body-colored females also dominated by generation 17 (98.31% at generation 17) (Figure 5B, C). By using DNA FISH, we found that this lineage carries yet another X chromosome (X**), where the Y-derived rDNA fragment was fused to the short arm of the X chromosome (Figure 5E, I-J). DNA FISH with Y short arm-specific satellite DNA (AATAAAC) confirmed that the rDNA fragment on the X** chromosome also originated from the Y chromosome (Figure 5J). The rapid expansion of pale body females in lineage 1E and lineage 2A occurred without any artificial selection, suggesting that fused X chromosomes carrying Y rDNA is strongly favored compared to carrying the Y chromosome. Indeed, we detected that the pale colored females have a slightly better fertility than dark colored females in lineage 1E (Figure 5K).

Taken together, the new karyotypes arising from the ‘*bb*^*158*^ stock’ provide further support that the full-length Y chromosome is disfavored in females: if the X chromosome loses sufficient rDNA copy number, and females become dependent on the Y chromosome as the source of rDNA, the karyotype quickly adapts to mitigate the burden of the Y chromosome in females.

## Discussion

Karyotype evolution is often associated with and speculated to be a cause of speciation (Ayala and Coluzzi, 2005; King, 1995; Lucek et al., 2022). However, the mechanisms that facilitate karyotype evolution are poorly understood. In this study, we provide examples of karyotype changes in *Drosophila* that involve rDNA. Combined with observations from other species (Page et al, 1996; Poot and Hochstenbach, 2021; Potapova and Gerton, 2019), we propose that rDNA instability may be a pervasive and universal causes of karyotype evolution.

We found that a *D. melanogaster* strain isolated from the Seychelles archipelago has a unique karyotype. This strain has lost most of the rDNA from their X chromosome, making XX females inviable. As a result, all females carry at least one Y chromosome as the source of rDNA. This strain also carries the truncated Y^S^ chromosome, which cannot support male fertility as it lacks a large portion of the long arm containing male fertility factors (e.g. axonemal dynein subunits). Our modeling suggests that the Y^S^ chromosome can be stably maintained in the population only if it has sufficiently smaller negative impact on female reproduction compared to the full-length Y chromosome. Indeed, our cytological analysis supports this idea that the Y^S^ chromosome is less deleterious than the full-length Y chromosome during oogenesis, in addition to the well-documented negative impact of Y chromosome heterochromatin on female longevity.

Strikingly, we found that a newly constructed strain that does not carry rDNA on the X chromosome (*bb*^*158*^/*bb*^*158*^ /Y females and *bb*^*158*^ /Y males) quickly developed new karyotypes. Two lineage developed new truncated Y chromosomes (Y^S^’ and Y^S^”) resembling the *D. mel*^Sey^ Y^S^ chromosome. In two other lineages, the fragment of Y chromosome containing rDNA fused to the X chromosome, generating new X* and X** chromosome fusions: Y-derived rDNA was fused to the long arm of X chromosome in X*, and to the short arm in X**. Females in these strains are now freed from the need of carrying a Y chromosome as the source of rDNA. Together, we observed five independent events of karyotype changes, where the absence of rDNA, combined with the negative impact of the Y chromosome on female fitness, led to karyotype changes.

In this study, we focused on the negative impact of the Y chromosome on female fitness as a driving force of karyotype changes. However, females needed to carry the Y chromosome due to loss of rDNA from the X chromosome. Given the universal instability of rDNA, it is tempting to speculate that the loss of rDNA loci, and adaptation to this loss, might serve as a general force for karyotype evolution.

Karyotype changes often occur by chromosome fusion/fission and centromere repositioning (Yoshida and Kitano, 2021), but these events can occur non-randomly (Kumon and Lampson, 2022). Interestingly, Robertsonian fusions, which are the most widespread karyotype changes that occur even within species (Poot and Hochstenbach, 2021), frequently involve the short arms of the telocentric chromosomes (Supplementary Figure 4) (Fel-Clair et al., 1998; Gravholt et al., 1992; Poot and Hochstenbach, 2021). In humans, Robertsonian fusions frequently occur between rDNA-bearing short arms, and rDNA loci have been speculated to play a role in inducing these translocations (Page et al, 1996; Poot and Hochstenbach, 2021; Potapova and Gerton, 2019). First, the homology between rDNA loci on non-homologous chromosomes may be sufficient for recombination, leading to the fusion of the chromosomes. Second, the well-established instability of an rDNA locus may play additional roles in triggering Robertsonian fusions: rDNA loci can lose their copy number, and sister chromatid recombination is known to mediate rDNA copy number recovery in yeast and *Drosophila* (Supplementary Figure 4) (Nelson et al., 2019). Accordingly, DNA breaks that are created in the process of sister chromatid recombination may further elevate the possibility of recombination between different rDNA loci.

Taken together, Robertsonian fusions in humans, as well as the examples of karyotype changes observed in *Drosophila* described in this study, point to an exciting possibility that rDNA instability may be a generalizable cause of karyotype evolution. The new karyotypes generated by rDNA instability may then be subject to selection for their fixation in the population, such as positive selection arising from a benefit (e.g., as a source of rDNA) or negative selection due to a disadvantage (e.g., toxic Y chromosome). In the cases described in the present study, a new karyotype appears to be selected to balance the necessity of rDNA against the toxicity of the Y chromosome on oogenesis. The fact that rDNA loci happen to be on sex chromosomes in *D. melanogaster* may add complexity to the selective force of karyotypes. In summary, we propose that rDNA instability may be a universal cause of karyotype evolution.

## Materials and Methods

### Fly husbandry and strains

Unless otherwise stated, all fly stocks were raised on standard Bloomington medium at room temperature (RT). The following D. melanogaster stocks were used: *yw*, Oregon-R, *Df(1)bb*^*158*^, *y1/Dp(1;Y)y+/C(1)\**, and *D. mel*^Sey^ (obtained from the University of California, San Diego Drosophila Stock Center, DSSC#14021-0231.123).

### Larval brain squash, DNA FISH, and microscopy

We adapted a simple FISH protocol against squashed chromosomes published by Larracuente and Ferree (2015) with small modifications. Briefly, male third instar wandering larvae were collected and brains were dissected in PBS. Larval brains were fixed in 25 ml of acetic acid: 4% formaldehyde in PBS (45%:55%) for 4 min on a clean Superfrost plus slide (Fisher Scientific) and the sample was manually squashed via thumb/stamp over coverslip, over sample, on top of the slide. The slide/coverslip/sample was immediately submerged in liquid nitrogen until it stopped boiling. Slides were removed from liquid nitrogen and coverslips were quickly flicked off the slide with a razor blade. Slides were then washed in 100% ethanol at room temperature for 5 min then dried in a dark, dust-free location. Hybridization was performed in 50% formamide, 10% dextran sulfate, 2 · SSC buffer, 0.5 mM of each probe, and up to 20 ml of diH2O. Hybridization buffer was added to the samples and covered with a coverslip. Slides were heated at 95°C for 5 min, cooled briefly, wrapped in parafilm, and incubated in a humid chamber in the dark overnight at room temperature. Coverslips were removed and slides were washed three times for 15 min in 0.1 · SSC, removed of excess buffer, and mounted in Vectashield mounting medium containing DAPI. Images were taken using an upright Leica Stellaris 8 or SP8 confocal microscope with a 63x oil immersion objective (NA = 1.4), and processed using Adobe Photoshop software. Images were modified solely for the purpose of clarity. Modified images were not quantified.

Detailed sequences of probes used are as follows:

IGS (part of 240 bp unit), 5’- AGTGAAAAATGTTGAAATATTCCCATATTCTCTAAGTATTATAGAGAAAAGCCATTTTAGTGA ATGGA-3’; (5’-AATAT-3’)_6_; (5’-AATAGAC-3’)_6_; (5’-AAAAC-3’)_6_; (5’-AAGAC-3’)_6_; (5’-AAGAGAG-3’)_5_; (5’-AATAC-3’)_6_; (5’-AATAAAC-3’)_6_; (5’-TAGA-3’)_8_

### Fertility Assay

Newly eclosed single males (or virgin females), were individually crossed to three yw virgin females (or newly eclosed males). After 7 days, each individual fly was subjected to genomic DNA extraction and karyotyping through ddPCR as described below. The number of adult flies eclosed from each vial was scored.

### Single-fly genomic DNA extraction

A single male fly was anesthetized under CO_2_ and transferred to a 200μL PCR tube on ice. A single mated female fly was dissected in PBS and only the carcass excluding ovaries and spermathecae was transferred to the PCR tube to minimize sperm DNA. 50μL squishing buffer (9.8mL dH2O, 100μL 1M Tris pH 7.5-8.0, 20μL 0.5M EDTA, and 50μL 5M NaCl) was mixed with 0.5μL of 20 mg/mL Proteinase K. The fly was smashed manually with a pipette tip before 50μL of the mixture was ejected into the PCR tube. The fly was further ground up with the buffer. The tube was placed in the thermocycler with the following program: 37°C for 30 minutes (digestion), 95°C for 2 minutes (heat-inactivate proteinase), and 4°C to hold. The Sample was spun down on table top centrifuge and 30μL of liquid was transferred to a clean tube to prepare for ddPCR karyotyping.

### Digital Droplet PCR (ddPCR) karyotyping

80 ng of DNA were used per 20 μL ddPCR reaction for both control gene reactions (RpL32 and Upf1) and Y chromosome reactions (Pp1-Y2 and PRY). Primers and probes for reactions are listed in Supplementary Table 1. ddPCR reactions were carried out according to the manufacturer’s (Bio Rad) protocol. In short, master mixes containing ddPCR Supermix for Probes (No dUTP) (Bio Rad), DNA samples, and primer / probe mixes were prepared in 0.2 mL Eppendorf tubes. ddPCR droplets were generated from samples using QX200 Droplet Generator (Bio Rad) and underwent complete PCR cycling on a C100 deep-well thermocycler (Bio Rad). Droplet fluorescence was read using the QX200 Droplet Reader (Bio Rad). Sample copy number was determined using Quantasoft software (Bio Rad). The total number of both Y and Y^S^ per genome was determined by calculating the ratio between Y linked genes and autosomal genes. A primer set for Pp1-Y2 was used to detect both Y and Y^S^ chromosomes, whereas a primer set for PRY was used to detect full length Y only. RpL32 on chromosome 3R was used as the denominator. Upf1 on the X chromosome was used as an additional control (chromosome 3:X should be either 1:1 in females or 2:1 in males). It should be noted that we frequently obtained values of Y:X close to 0.8 (instead of 1.0) when there is one copy of each, presumably due to the different efficiencies of PCR primers. 0.8 was considered as Y:X = 1:1.

### Immunofluorescence on whole-mount ovaries

Immunofluorescence staining was performed as described previously (Cheng et al., 2008), with slight modifications for single fly. Briefly, tissue from individual female flies were dissected in PBS (while the carcasses were subject to genomic DNA extraction), transferred to 4% formaldehyde in PBS and fixed for 30 min. Tissues were then washed in PBS-T (PBS containing 0.1% Triton-X) for at least 60 min, dehydrated in 50%:50% methanol:PBS-T for 10 min, and two times in 100% methanol for 10 min before frozen in methanol at -20°C until staining. When ready to stain, tissues were rehydrated in 50%:50% methanol:PBS-T and then two times of PBS-T for 10 min each, followed by incubation with primary antibody in 3% bovine serum albumin (BSA) in PBS-T at 4°C overnight. Samples were washed for 60 min (six 10 min washes) in PBS-T, incubated with secondary antibody in 3% BSA in PBS-T at 4°C overnight, washed as above, and mounted in VECTASHIELD with DAPI (Vector Labs). The following primary antibodies were used: rat anti-vasa (1:20; DSHB; developed by A. Spradling), and mouse anti-Fasciclin III (1:200; DSHB; developed by C. Goodman). Alexa Fluor-conjugated secondary antibodies (Abcam) were used with a dilution of 1:200. Images were taken using a Leica Stellaris 8 confocal microscope with 10x dry objectives (NA = 0.4) and 63x oil-immersion objectives (NA = 1.4). Images were processed using Adobe Photoshop software.

### Sex chromosome frequency simulations

There are several assumptions in our simulation. First, in females, when there are two X chromosomes and one Y or Y^S^ chromosome, two X chromosomes pair and segregate whereas the Y or Y^S^ chromosome accompanies one of the two X chromosomes in meiosis I. For example, XXY females are assumed to produce 50% X eggs and 50% XY eggs. Based on our observation, frequencies of X nondisjunction (i.e., production of XX eggs) in XXY and XXY^S^ are comparable to XX females which are assumed to produce only X eggs, ignoring infrequent X nondisjunction events. Second, two X chromosomes pair and segregate and so do two Y chromosomes since in *Drosophila* females pairing is based on homology (Hughes et al., 2018). For example, XXYY^S^ females are assumed to produce 50% XY eggs and 50% XY^S^ eggs. Third, sex chromosome trivalents of males segregate randomly in meiosis I. For example, in meiosis I of XYY^S^ males, one third of the time X and YY^S^ segregate, XY and Y^S^ segregate, and XY^S^ and Y segregate. This assumption is based on the empirical observation that in meiosis I of both XYY and XYY^S^ males, any one of the three sex chromosomes segregates from the other two around ⅓ of the time. Likewise, sex chromosome quadrivalents (e.g., XYY^S^Y^S^) of males are assumed to segregate randomly in meiosis I. Possible gamete genotypes for each parent sex chromosome genotype is summarized in Supplementary Figure 2A.

Fitness values for each parental genotype are summarized in Supplementary Figure 2A. As the Seychelles X chromosomes lack rDNA, XX fitness is set to zero. Values of *s* and *y* define the female cost of Y^S^ and Y chromosomes when present as two copies, respectively (“Y^S^ Cost” and “Y Cost” in Figure 4). Values of *s* and *y* vary between 0 and 1. The value of *h* defines the cost of having one Y^S^ chromosome (XXY^S^), relative to the cost of having two Y^S^ chromosomes (XXY^S^Y^S^) in females. The value of *h’* defines the cost of having one Y chromosome (XXY), relative to the cost of having two Y chromosomes (XXYY) in females. The value of *h”* defines whether YY^S^ heterozygous fitness is more similar to Y^S^Y^S^ homozygous fitness (*h”*=0) or YY homozygous fitness (*h”*=1) in females. Values of *h, h’* and *h”* vary between 0 and 1, and we assumed *h*=*h’*=*h’’*=0.5, which best describes our observed female fitness in Figure 3.

Fitness values of males are binary (0 or 1) except for XYYY^S^ and XYY^S^Y^S^. As the Seychelles Y^S^ chromosomes lack essential male fertility factors like axonemal dynein, any male genotype that carries only Y^S^ chromosomes is set to zero. Having up to two Y or Y^S^ chromosomes does not impact male fertility (Figure 2I). However, having three or more than three Y or Y^S^ chromosomes is likely deleterious, as three Y or Y^S^ individuals are underrepresented in the Seychelles population (Figure 2F, see also main text for deleterious effects of heterochromatin). XYYY individuals are never observed, so their fitness is set to zero. XYYY^S^ and XYY^S^Y^S^ fitness values are set to 0.1, which best describes our observed number of three Y or Y^S^ individuals (Figure 2F and Supplementary Figure 2B).

For each individual (at the fertilized egg stage) in each generation, sex chromosome genotype is randomly determined by the frequencies of gamete genotypes from the preceding generation. We used the observed frequencies (Figure 2E-F) to simulate the first generation. The individual genotype of the fertilized egg is then used to calculate the “individual contribution to the gamete pool”. The individual contribution to the gamete pool is calculated by the fitness value (ranges between 0 and 1) multiplied by the frequency of gamete genotype for each parent genotype. For example, a XY male individual produces 1 (fitness value) x 0.5 (frequency of X sperm) = 0.5 X sperm and 1 (fitness value) x 0.5 (frequency of Y sperm) = 0.5 Y sperm, whereas a XX female individual produces 0 (fitness value) x 1 (frequency of X eggs) = 0 X eggs. For the population size of 100 individuals, the individual contribution to the gamete pool is summed up and used to calculate the frequencies of gamete genotype in this generation, which is then used to simulate the parental genotypes in the following generation. The custom codes used to simulate sex chromosome frequenceis are available on the GitHub (https://github.com/TomoKumon/).

## Acknowledgments

We thank the Bloomington Drosophila Stock Center, UCSD Drosophila stock center, Developmental Studies Hybridoma Bank for reagents. We thank the Yamashita lab and the Lehmann lab members for insightful discussions, the Yamashita lab members for the comments on the manuscript. The research was supported by the Howard Hughes Medical Institute and Whitehead Institute for Biomedical Research.

## Notes

### Competing Interest Statement

The authors have declared no competing interest.

