## supplementary material for "ribosomal DNA instability as a potential cause of karyotype evolution"

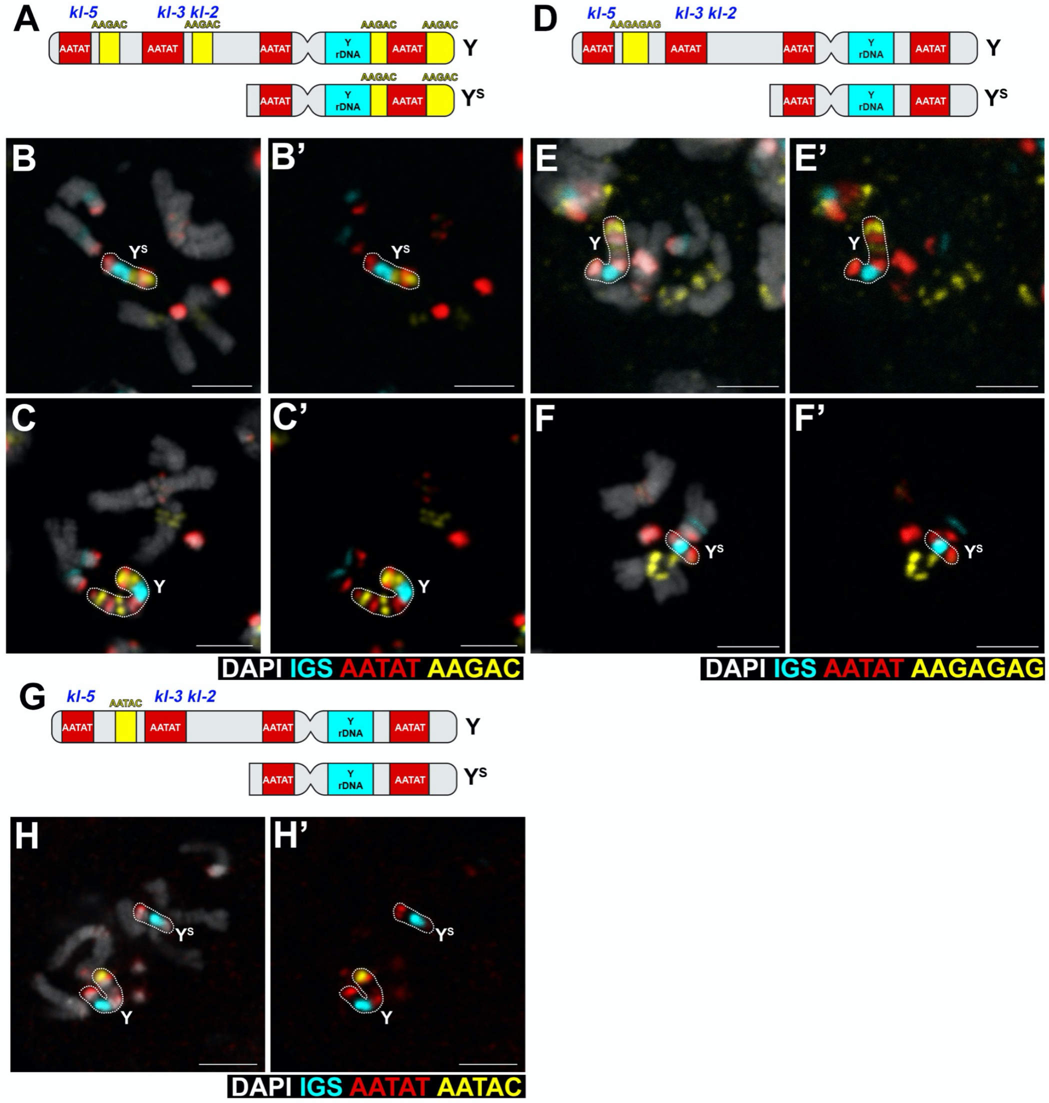


**Supplementary Figure 1. Panel of satellite DNAs used to differentiate Y and $Y^S$ chromosomes.**

A-C) DNA FISH on larval neuroblast mitotic chromosome spread showing AAGAC satellite is present on Y but not $Y^S$. Approximate locations of kl-5, kl-3, kl-2 are shown. Bar: 3µm in all panels.

D-F) DNA FISH on larval neuroblast mitotic chromosome spread showing AAGAGAG satellite is present on Y but not $Y^S$.

G-H) DNA FISH on larval neuroblast mitotic chromosome spread showing AATAC satellite is present on Y but not $Y^S$.

**A**

| Female Genotype | Fitness | Egg Genotype (Frequency) |
|---|---|---|
| XX | 0 | — |
| $XXY^S$ | 1–*hs* | X (1/2), $XY^S$ (1/2) |
| XXY | 1–*h'y* | X (1/2), XY (1/2) |
| $XXY^SY^S$ | 1–*s* | $XY^S$ (1) |
| $XXYY^S$ | (1–*s*)–*h''*(*y*–*s*) | XY (1/2), $XY^S$ (1/2) |
| XXYY | 1–*y* | XY (1) |

| Male Genotype | Fitness | Sperm Genotype (Frequency) |
|---|---|---|
| XY | 1 | X (1/2), Y (1/2) |
| $XY^S$ | 0 | — |
| XYY | 1 | X (1/6), YY (1/6), XY (2/6), Y (2/6) |
| $XYY^S$ | 1 | X (1/6), $YY^S$ (1/6), XY (1/6), $Y^S$ (1/6), $XY^S$ (1/6), Y (1/6) |
| $XY^SY^S$ | 0 | — |
| XYYY | 0 | — |
| $XYYY^S$ | 0.1 | XY (2/6), $YY^S$ (2/6), $XY^S$ (1/6), YY (1/6) |
| $XYY^SY^S$ | 0.1 | XY (1/6), $Y^SY^S$ (1/6), $XY^S$ (2/6), $YY^S$ (2/6) |
| $XY^SY^SY^S$ | 0 | — |

**B**

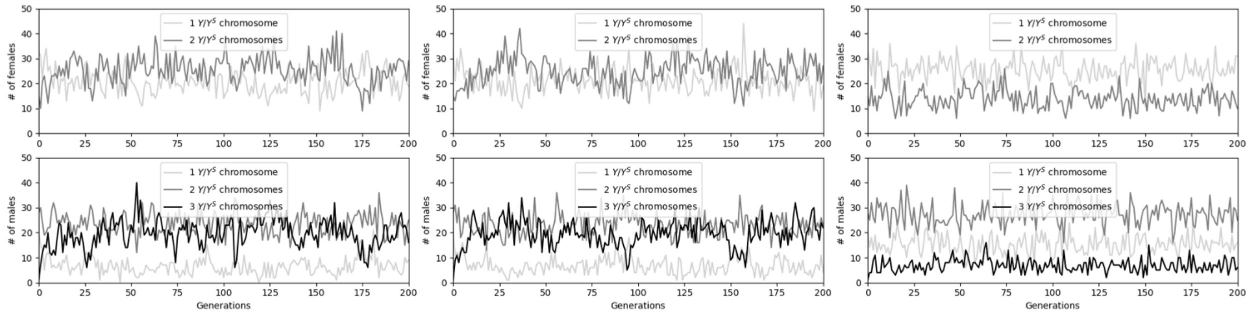


**Supplementary Figure 2. The simulation using various parameters.**

A) Relative fitness values of each parental genotype. For each parental genotype, possible gamete genotypes and their frequencies are shown. See Method.

B) The three panels simulate the number of males and females over two hundred generations with different male fitness values. "1 $Y/Y^S$ chromosome" individuals carry one Y or $Y^S$ chromosome, "2 $Y/Y^S$ chromosomes" carry two, and "3 $Y/Y^S$ chromosomes" carry three. The female fitness values are the same for all three simulations ($y$=0.90, $s$=0.45, $h$=$h'$=$h''$=0.50). The left panel assumes XYYY, $XYYY^S$, and $XYY^SY^S$ are all fertile (fitness value = 1), whereas the middle panel assumes XYYY is infertile (fitness value = 0) but $XYYY^S$ and $XYY^SY^S$ are fertile (fitness value = 1). In both cases, the number of males that carry three Y or $Y^S$ chromosomes is overrepresented than our observation in Figure 2F. The right panel assumes XYYY is infertile (fitness value = 0) and $XYYY^S$ and $XYY^SY^S$ are subfertile (fitness value = 0.1). Since this parameter set best describes the observed male frequencies, the male fitness value of 0.1 is assumed for $XYYY^S$ and $XYY^SY^S$. See Method.

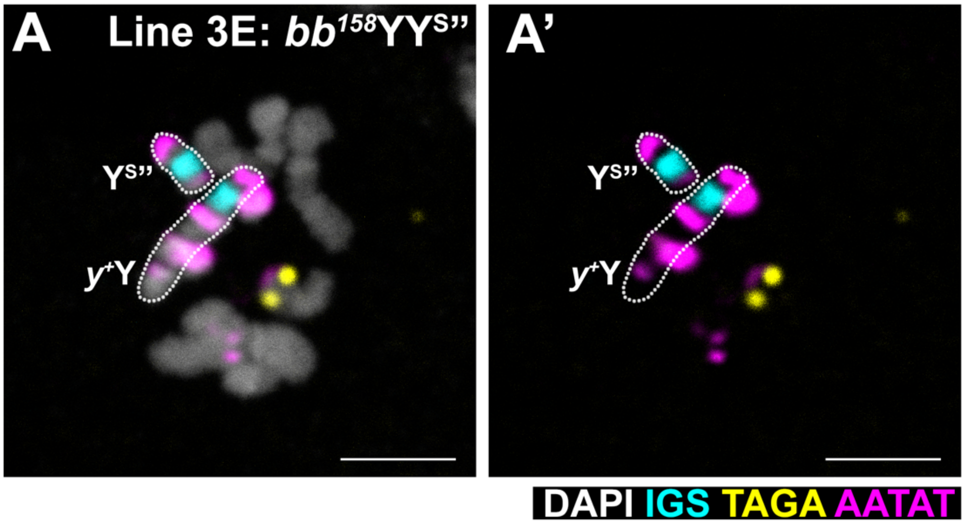


**Supplementary Figure 3.** DNA FISH on a larval neuroblast mitotic chromosome spread from a *bb*$^{158}$/*y*$^{+}$Y/Y$^{S}$*”* male. Bar: 3 µm.

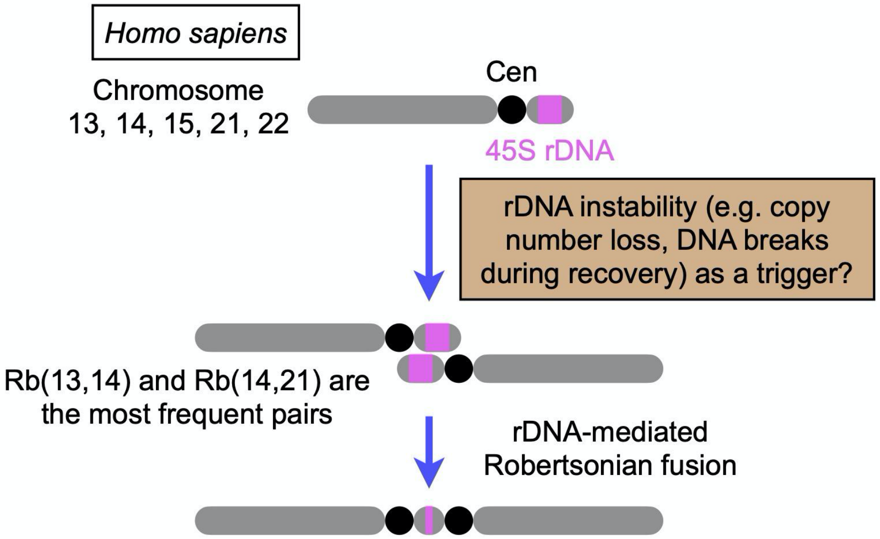


**Supplementary Figure 4: rDNA instability can explain frequent Robertsonian fusion events in human chromosomes.**
Human chromosomes are mostly metacentric except for chromosomes with 45S rDNA, where they are found on the short arm (Potapova and Gerton, 2019). In human, Robertsonian fusion events are frequently found in chromosomes with the 45S rDNA (Page et al, 1996; Poot and Hochstenbach, 2021).

| Oligo Name | Type | Target | Sequence | Modifications |
|---|---|---|---|---|
| dd-RpL32 F | Primer | RpL32 | GCTTCAAGGGACAGTATCTG | |
| dd-RpL32 R | Primer | RpL32 | AACGCGGTTCTGCATGAG | |
| dd-RpL32 Probe | Probe | RpL32 | ATGCCCAACATCGGTTAC | 5' HEX AND 3' Iowa Black FQ |
| dd-Upf1 F | Primer | Upf1 | CACACTTTATGTCCACCATTATTG | |
| dd-Upf1 R | Primer | Upf1 | GAGTTTCCGTAGGGACCAC | |
| dd-Upf1 Probe | Probe | Upf1 | CCG TAA CCG CCA CTG CGG T | 5' 6-FAM AND 3' Iowa Black FQ |
| dd-Pp1-Y2 F | Primer | Pp1-Y2 | GTCGCAACCAATGCTCC | |
| dd-Pp1-Y2 R | Primer | Pp1-Y2 | GGTAATTGGACGCTGGTGG | |
| dd-Pp1-Y2 Probe | Probe | Pp1-Y2 | CAGCCTCAATAGGTCAGTAAACTGAC | 5' 6-FAM AND 3' Iowa Black FQ |
| dd-PRY F | Primer | PRY | CCACAAACAACAGTCCAGCTCG | |
| dd-PRY R | Primer | PRY | CCATGTCATCAAGTGGTTCCAAGG | |
| dd-PRY Probe | Probe | PRY | CTCACCTCGTTCGCCTTACAAAGAGGGTT | 5' HEX AND 3' Iowa Black FQ |

**Supplementary Table 1: List of oligos used in ddPCR assays.**

Primers and Probe for RpL32 and Upf1 are from Nelson et al. 2021
